# Accelerated cell cycles enable organ regeneration under developmental time constraints in the *Drosophila* hindgut

**DOI:** 10.1101/2020.02.17.953075

**Authors:** Erez Cohen, Donald T. Fox

## Abstract

Individual organ development must be temporally coordinated with development of the rest of the organism. As a result, cell division in a developing organ occurs on a relatively fixed time scale. Despite this, many developing organs can regenerate cells lost to injury. How organs regenerate within the time constraints of organism development remains unclear. Here, we show the developing *Drosophila* hindgut regenerates by accelerating the mitotic cell cycle. This process requires JAK/STAT signaling and is achieved by decreasing G1 length during the normal period of developmental mitoses. Mitotic capacity is then terminated by the steroid hormone ecdysone receptor. This receptor activates a hindgut-specific enhancer of *fizzy-related*, a negative regulator of mitotic cyclins. We further identify the Sox transcription factor *Dichaete* as an important negative regulator of injury-induced mitotic cycles. Our findings reveal how mitotic cell cycle entry mechanisms can be adapted to accomplish developmental organ regeneration.

## Introduction

Development is subject to exquisite temporal regulation (Nüsslein-Volhard and Wieschaus, 1980, reviewed in Lagha et al., 2012). This control synchronizes development of the organ with development of the organism. For example, during *Drosophila* metamorphosis, systemic hormonal signals simultaneously time the growth of specific tissues and also the molting of the whole animal (Karim and Thummel 1992; reviewed in Riddiford *et al*. 2000; Kozlova and Thummel 2003).

In spite of the tight temporal regulation of development, many developing tissues have a striking capacity to regenerate after injury. Examples of such developmental regeneration across the animal kingdom include the *Xenopus* tadpole tail, the *Drosophila* imaginal discs, and the myocardium and digit tips of developing mammals (Halme et al., 2010; Illingworth, 1974; Porrello et al., 2011; Slack et al., 2004; Smith-Bolton et al., 2009). If developing tissues are unable to repair an injury, long-term tissue abnormalities may arise. For example, pediatric traumatic brain injury causes detrimental neural developmental abnormalities (Imms et al., 2019; Taylor et al., 2017) and reduced brain function (Ganesalingam et al., 2011; Lindsey et al., 2019). In bone tissue, deformities can occur after unresolved pediatric facial fractures (<5 years) (Singh and Bartlett, 2004; Wheeler and Phillips, 2011). Thus, while development is subject to temporal constraints, the capacity for organ repair and regeneration plays important roles across evolution.

In *Drosophila,* one paradigm of developmental injury repair involves extending the time needed to complete organism development. *Drosophila* larval imaginal tissues consist of progenitor cells that make up the organs of the adult fly. Imaginal discs undergo compensatory cell divisions to regenerate disc-derived tissues following tissue injury (reviewed in Hariharan and Serras 2017). In the wing imaginal disc, these extra regenerative divisions do not occur within normal organismal developmental timing. Rather, wing injury at 2^nd^ and early 3^rd^ instar larval stages releases systemic cues that activate a regeneration checkpoint. This checkpoint delays animal development, allowing the wing disc extra time to both repair and properly develop (Halme et al., 2010; Hussey et al., 1927; Simpson et al., 1980; Smith-Bolton et al., 2009). Once development passes the period of checkpoint activation, the wing disc can no longer regenerate (Halme et al., 2010; Smith-Bolton et al., 2009). Currently, existing models in *Drosophila* have been unable to study how organs might complete tissue repair within developmental time constraints.

The *Drosophila* hindgut consists of three main segments-the pylorus, ileum, and rectum. These segments are present in both the larval and adult gut. During metamorphosis, the normally developing hindgut undergoes a regeneration of sorts. As documented by Robertson over 80 years ago (Robertson, 1936), the larval ileum undergoes histolysis, and is replaced by regenerative activity from adult hindgut precursors. Cell division from the larval pylorus, which occurs during metamorphosis, is the source of the expanded adult pylorus, and also the new adult ileum (Cohen et al., 2020; Fox and Spradling, 2009; Sawyer et al., 2017; Takashima et al., 2008; Yang and Deng, 2018). This development is not disrupted by a severe, acute injury to the wandering 3^rd^ instar larval pylorus. Our previous lineage labeling suggested that pyloric cells remaining after injury undergo additional cell divisions (Cohen et al., 2018). Nevertheless, the capacity of the pylorus to respond to injury by regenerative cell divisions is lost by adulthood. Unlike the larval pylorus, injury to the adult pylorus results in endocycles, a cell cycle in which cells replicate DNA without mitosis, leading to polyploidization and hypertrophy (Cohen et al., 2018; Fox and Duronio, 2013; Fox and Spradling, 2009; Losick et al., 2013; Sawyer et al., 2017). This switch in injury response from larval mitosis to adult endocycles is regulated by expression of the Anaphase Promoting Complex/Cyclosome (APC/C) activator Fizzy-related (Fzr), also known as Cdh1 (Cohen et al., 2018). The developmental activation of *fzr*, which occurs irrespective of injury, implies a role for developmental signals in switching injury responses and limiting pyloric mitotic capacity. The hindgut pylorus can therefore be studied as a model to uncover how developmental signals can alter tissue injury responses.

Here, we report that unlike the wing imaginal disc, the hindgut can regenerate without delaying whole animal development. Instead, hindgut regeneration after injury occurs through an acceleration of mitotic cell cycle rate. This enables the hindgut to undergo additional mitotic cell cycles within the normal time frame of mitosis during hindgut development. This cell cycle acceleration occurs by reducing G1 phase length, and requires Unpaired3, a JAK/STAT pathway cytokine. We further show that the time window in which the rapid regenerative mitotic cycles occur is terminated by developmental activation of a *fzr* enhancer. This enhancer is activated by the steroid hormone Ecdysone Receptor (EcR). Additionally, this *fzr* enhancer contains binding sites for the Sox-domain-containing transcription factor *Dichaete,* which (like *fzr*) negatively regulates injury-induced hindgut mitosis. Our findings reveal that the hindgut pylorus is a model to uncover control of mitotic organ regeneration under developmental time constraints.

## Results

### The *Drosophila* hindgut can regenerate without delaying metamorphosis

To understand how organ regeneration is accomplished in the hindgut despite possible time constraints imposed by development (metamorphosis), we considered two possible models. In the first model, hindgut injury leads to an organism-wide developmental delay (**Fig1A, Model1**) until injury is repaired, allowing time for additional regenerative cell cycles. In a second alternative model, acute injury does not delay organism development, assayed by pupation onset (**Fig1A, Model2**). In this model, additional compensatory mitotic cell cycles would have to occur within a normal developmental time frame. To distinguish between these two models, we acutely injured the larval hindgut and assessed whole-animal development progression (**Methods**). For tissue injury, we used the temporally and spatially regulated Gal4-UAS expression system, driven by a hindgut-specific enhancer of the *brachyenteron (byn)* gene to induce the apoptotic genes *head involution defective* (*hid*) and *reaper* (*rpr*) (Cohen et al., 2018; Fox and Spradling, 2009; Losick et al., 2013; Sawyer et al., 2017; Smith-Bolton et al., 2009). We induced injury at either the 2^nd^ or early 3^rd^ larval instar (L2-L3) stages or at the wandering third instar (hereafter L3W) stage. To assess whether whole animal developmental progression timing changes following injury, we measured the time of pupation onset in animals with or without *hid* and *rpr* transgenes.

**Figure1.**
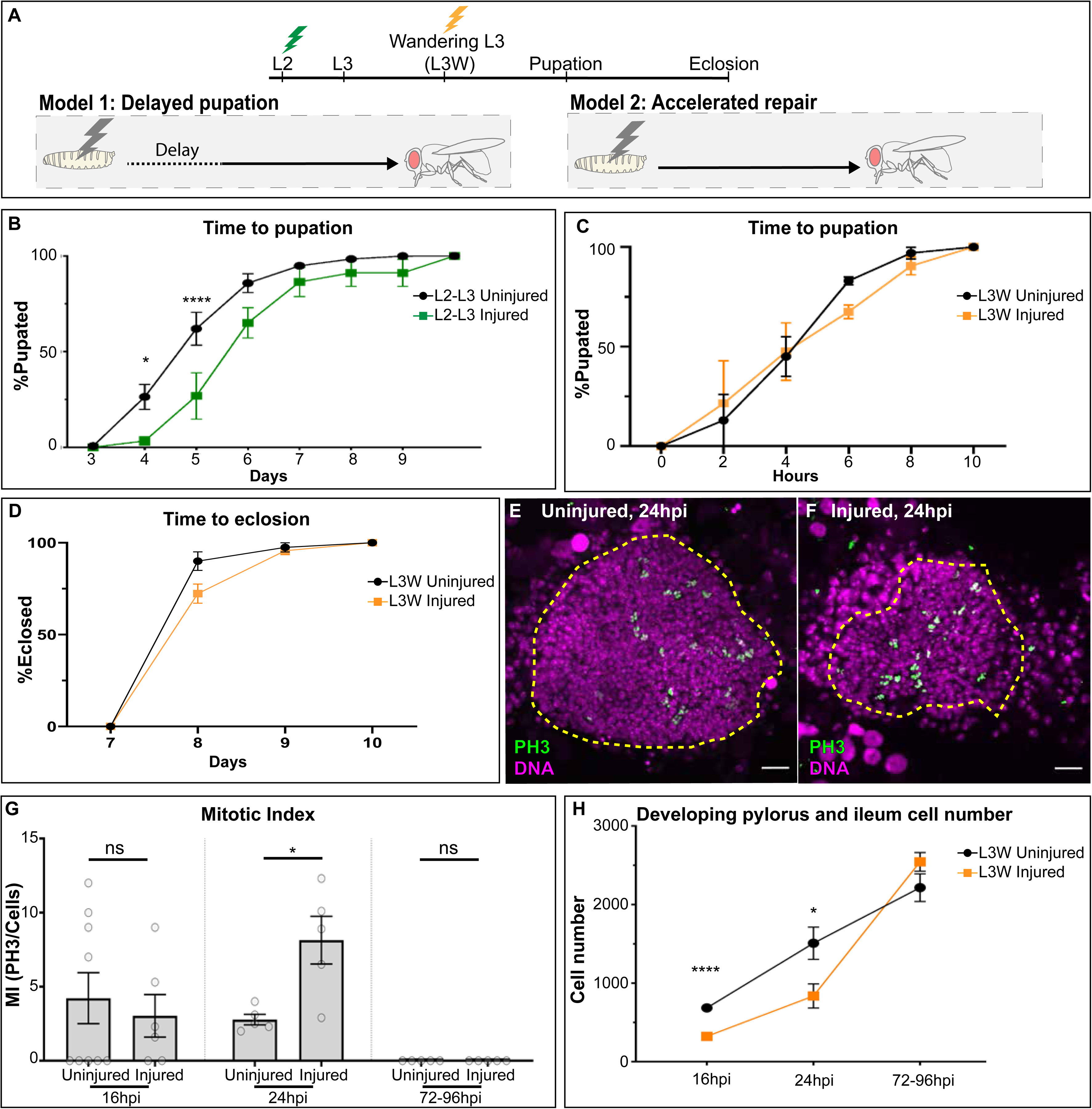
The injured wandering 3^rd^ instar hindgut regenerates by increasing mitosis without delaying organismal development. (**A**) Schematic illustrating two models for regeneration under developmental time constraint. Lightning bolts represent injury induction. (**B**) Time from 3^rd^ instar larva to pupation onset in L2-early L3 animals with injured hindguts or uninjured controls. Data represent mean ± SEM, N≥5 animals per condition, at least two replicates. Two-way ANOVA (interaction p<0.05), Sidak’s multiple comparisons reported in graph. (**C-D**) Time from L3W to (**C**) pupation onset or (**D**) the adult stage in L3W animals with injured hindguts or uninjured controls. Data represent mean ± SEM, N≥5 animals per condition, at least two replicates. Two-way ANOVA (interaction and condition not significant). (**E-F**) Mitotic figures at 24 hours post injury (hpi) in the (**E**) uninjured pylorus or (**F**) pylorus injured at L3W. Phospho-HistoneH3 (green), and nuclei (DAPI, magenta) are marked. Yellow hashed line outlines developing pylorus, derived from *byn* expression(**G**) Quantification of pyloric mitotic index (MI, %mitotic cells) in the presence or absence of injury at L3W. Data represent mean ± SEM, N≥5 animals per condition, at least two replicates. Unpaired two-tailed t-tests. (**H**) Cell number in developing pylori in the presence or absence of injury at L3W. Data represent mean ± SEM, N≥5 animals per condition, at least two replicates. Unpaired two-tailed t-tests reported in graph. Analysis of recovery slope using ANCOVA identifies p<0.05 for 24-72hr slope. Scale bars 20µm.

As previous experiments found a whole-organism developmental delay when the wing is injured at the L2-early L3 stage, we first injured the hindgut at this stage. Similar to the wing, acute L2-early L3 hindgut injury leads to a significant developmental delay (**Fig1B**). The delay in pupation onset is approximately 24 hours, similar to the delay observed following *rpr-*induced injury of the L2-early L3 wing imaginal disc (Halme et al., 2010). As with our previous experiments using L3W injury, L2-early L3 hindgut injury leads to wholescale organ regeneration (data not shown). In addition to injuring at the L2-early L3 stage, we previously established that, unlike in the wing, the hindgut is capable of whole organ regeneration when injury is induced at the L3W stage, shortly before pupation onset (Cohen et al., 2018). We therefore tested whether injury to the L3W hindgut results in organismal development delay. In contrast to injury at the L2-early L3 stage, injury to the L3W hindgut does not lead to organism-wide delay in pupation onset (**Fig1C**). However, it remained possible that injury to the L3W hindgut causes a developmental delay during pupation. We tested for any delay in pupation progression by measuring time of pupation (L3W to eclosion). We find there is no delay in fly eclosion following L3W hindgut injury (**Fig1D**). These results indicate that injury to different stages of the larval hindgut results in different developmental responses. At L2-early L3, hindgut injury initiates a developmental delay, while at L3W, hindgut injury leads to whole organ regeneration in concert with animal development.

We next looked at the cellular level to begin to understand how the injured L3W hindgut is able to fully regenerate without delaying development. As the pylorus is the site of hindgut regenerative activity, we closely examined this region. We assayed cell death, cell number, and the frequency of pyloric cells with the mitotic marker Phospho-Histone H3 (PH3, **Methods**) during metamorphosis in animals injured at L3W. At 16 hours post-injury (hpi), we observe widespread apoptosis and reduced cell number (**FigS1A-A’, Fig1H,** 16hpi). However, over the course of metamorphosis, injured animals display an accelerated rate of cell number increase, along with a higher mitotic index at 24hpi (**Fig1E-H**). Consistent with our results that development is not delayed, we do not see a broadening of the mitotic window as observed by examining the mitotic index of late (72-96hpi) pupal pylori (**Fig1G, FigS1B-D**). Further, cell number is restored within the normal pyloric developmental timeframe (**Fig1H, FigS1C-D**).

We next identified the number of additional mitotic cell cycles that the L3W pylorus must undergo following injury to produce the expanded adult pylorus and also the adult ileum. In the absence of injury, our previous lineage tracing experiments identified that L3W pyloric cells destined to produce the adult pylorus undergo approximately 2-3 divisions (yielding 5-6 cell clones in adults), whereas pyloric cells destined to generate the adult ileum undergo one division to make up the adult ileum (yielding 2 cell clones in adults) (Cohen et al., 2018; Fox and Spradling, 2009; Sawyer et al., 2017). In the presence of injury, our lineage tracing following L3W injury indicated a 3-fold increase in recovered adult pyloric clone size (from 5-6 cells to ∼16 cells) and a doubling of ileal clone size (from 2 cells to 4 cells, Cohen et al., 2018). Given our prior counts of adult pyloric and ileal cells (Cohen et al., 2018; Fox and Spradling, 2009; Sawyer et al., 2017) and our lineage observations, we conclude that our injury protocol causes adult hindgut progenitors, on average, to undergo 2-3 additional mitotic cell cycles. Consistent with this finding, our data identifies a 2-3x fold increase in mitotic index at 24hpi (2.8±0.4% vs 8.1±1.6%). These data also argue that the increased mitotic index after injury does not simply reflect a prolonged progression through M-phase. Additionally, through revisiting our previous lineage tracing data (Cohen et al., 2018), we also ruled out the model that injury activates a dormant, injury-responsive pool of mitotic pyloric cells. Specifically, we find no change in clone number after injury (**FigS1E**). Taken together, our results pinpoint a timeframe within pupal development when pyloric cells undergo additional divisions to compensate for acute injury.

### The JAK/STAT cytokine Unpaired3 and a shortened G1 phase accelerates the cell cycle during developmental hindgut regeneration

We next sought to identify regulators of the increased cell divisions that regenerate the hindgut under developmental time constraints. We focused on activation of the JAK/STAT pathway ligand Unpaired3 (Upd3, an IL6-like ligand). Previous research identified a role for *upd3* in injury-induced cell cycle entry in various *Drosophila* tissues including hemocytes (Pastor-Pareja et al., 2008), adult midgut intestinal stem cells (Biteau et al., 2008; Jiang et al., 2009; Osman et al., 2012) and adult hindgut pyloric cells (Sawyer et al., 2017). To specifically test *upd3* function, we examined animals containing an X-chromosome null mutation of *upd3 (upd3^Δ^)*. Consistent with *upd3* functioning primarily in stress responses, *upd3^Δ^* homozygous animals are viable, fertile, and do not show any major animal-wide developmental defects (Osman et al., 2012). We first tested the requirement of *upd*3 in hindgut development by assessing the morphology of adult hindguts from animals hemizygous for the mutant allele *upd3^Δ^.* We do not observe morphological defects or cell loss in *upd3^Δ^* hindguts, indicating that *upd3* is not essential for hindgut development (**Fig2A, D**). In sharp contrast to the recovery observed after L3W hindgut injury in wild-type animals (**Fig1H, 72**hpi), acute L3W hindgut injury in *upd3^Δ^* animals leads to gross hindgut abnormalities (**Fig2B, D**). 100% of injured *upd3^Δ^* animals die within 3 days following eclosion. Notably, hindgut abnormalities and animal lethality also occur following injury in heterozygous *upd3^Δ^* females (**Fig2C, D**). Our data suggest that correct levels of *upd3* are essential for the L3W hindgut injury response, but not for normal hindgut metamorphosis.

**Figure2.**
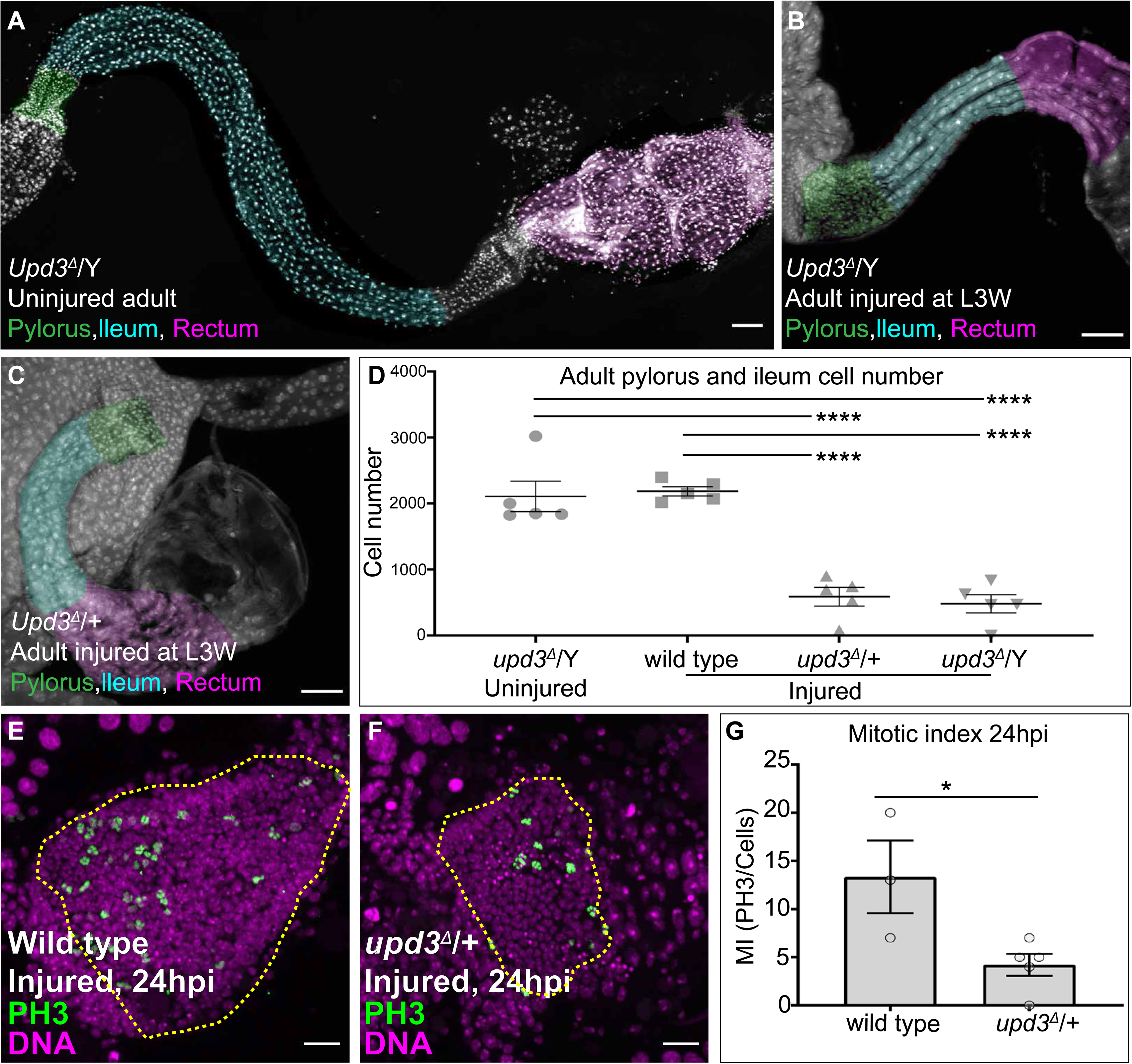
The JAK/STAT ligand Upd3 is essential for regenerative divisions. **(A-C)** Hindgut morphology in adult animals. (**A**) uninjured hemizygous *upd3^Δ^*, (**B**) injured hemizygous *upd3^Δ^,* and (**C**) heterozygous *upd3^Δ^* animals recovered from injury at L3W. Nuclei (DAPI, white) and false coloring identifies the tissues of the adult hindgut: Pylorus (green), Ileum (cyan) and Rectum (magenta). Pairwise stitching was performed to represent the entire hindgut in a single image. (**D**) Quantification of adult pylorus and ileal cell number (combined) after injury to the wandering L3 pylorus in wild-type and *upd3* mutants. Data represent mean ± SEM, N≥5 animals per condition, at least two replicates. ANOVA, Tukey multiple comparisons test. (**E-F**) Mitotic figures 24 hours post injury of wandering L3 pylorus in (**E**) wild-type and (**F**) *upd3* heterozygous animals. Phospho-HistoneH3 (green), and nuclei (DAPI, white) are marked. Yellow hashed line outlines developing pylorus, derived by *byn* expression. (**G**) Quantification of pyloric mitotic index (MI, %mitotic cells) 24 hours post wandering L3 injury in wild-type and *upd3* heterozygous animals. Data represent mean ± SEM. N=5 animals for heterozygous *upd3 ^Δ^* condition and N=3 for wild-type condition, at least two replicates. For additional wild-type control animal data see Figure1 panel I. Unpaired two-tailed t-test. Scale bars (**A-C**) 50µm, (**E-F**) 20µm

Next, we tested whether *upd3* is required in pyloric cells to increase mitotic index following acute L3W injury. The mitotic index of injured heterozygous *upd3^Δ^* females at 24hpi does not increase in response to injury and resembles wild-type, uninjured animals (**Fig2E-G, Fig2G:***upd3*/+ **vs**. **Fig1G:**24hpi uninjured). Together, our results support a role for the *upd3* cytokine in responding to hindgut injury by increasing pyloric cell mitotic activity during pupation. This increase allows for additional, injury-mediated cell divisions within developmental time constraints.

Given our findings that Upd3-mediated signaling increases the number of pyloric cell mitotic divisions after L3W hindgut injury, we next examined whether any cell cycle phase is shortened to facilitate these extra divisions. We assessed progression of the pyloric cell cycle using the Fly-FUCCI system with and without injury. FUCCI reporters utilize a set of markers (GFP-E2F11-230 and RFP-CycB1-266) to distinguish cells at G1, S and G2/M stages (Zielke et al., 2014). Combining our DEMISE injury system (Cohen et al., 2018) with Fly-FUCCI allowed us to simultaneously injure animals using FLP recombinase-mediated activation of *rpr* while also expressing FUCCI reporters in cells adjacent to apoptotic cells in the developing hindgut (**Fig3A**). Consequently, we were able to assay the distribution of cell cycle stages across development in the presence or absence of injury (**Methods**). Upon injury, we find a reduction in the percentage of pyloric cells in G1 at both 16 and 24hpi as seen by increased GFP-E2F11-230 positive, RFP-CycB1-266 negative cells (**Fig3B-F**). By completion of hindgut development and recovery (as determined in **Fig1H and FigS1B**), all cells are at the G1/G0 stage irrespective of previously induced injury (**Fig3F**). Given the persistence of the GFP-E2F11-230 marker in nearly all pyloric cells, we did not distinguish between S and G2 stages. Compared to developing uninjured animals, injured pylori show an increase in S/G2 cells immediately following injury (**Fig3G**). Together, our FUCCI results indicate that injury causes a doubling in pyloric cells leaving G1 to enter S/G2 stages (**Fig3F-G,** 16hpi), followed by a doubling in mitotic index soon after (**Fig1G,** 24hpi). Therefore, by accelerating pyloric cell cycle speed through shortening G1, the injured L3W hindgut achieves compensatory mitotic cycling within normal developmental time constraints.

**Figure3.**
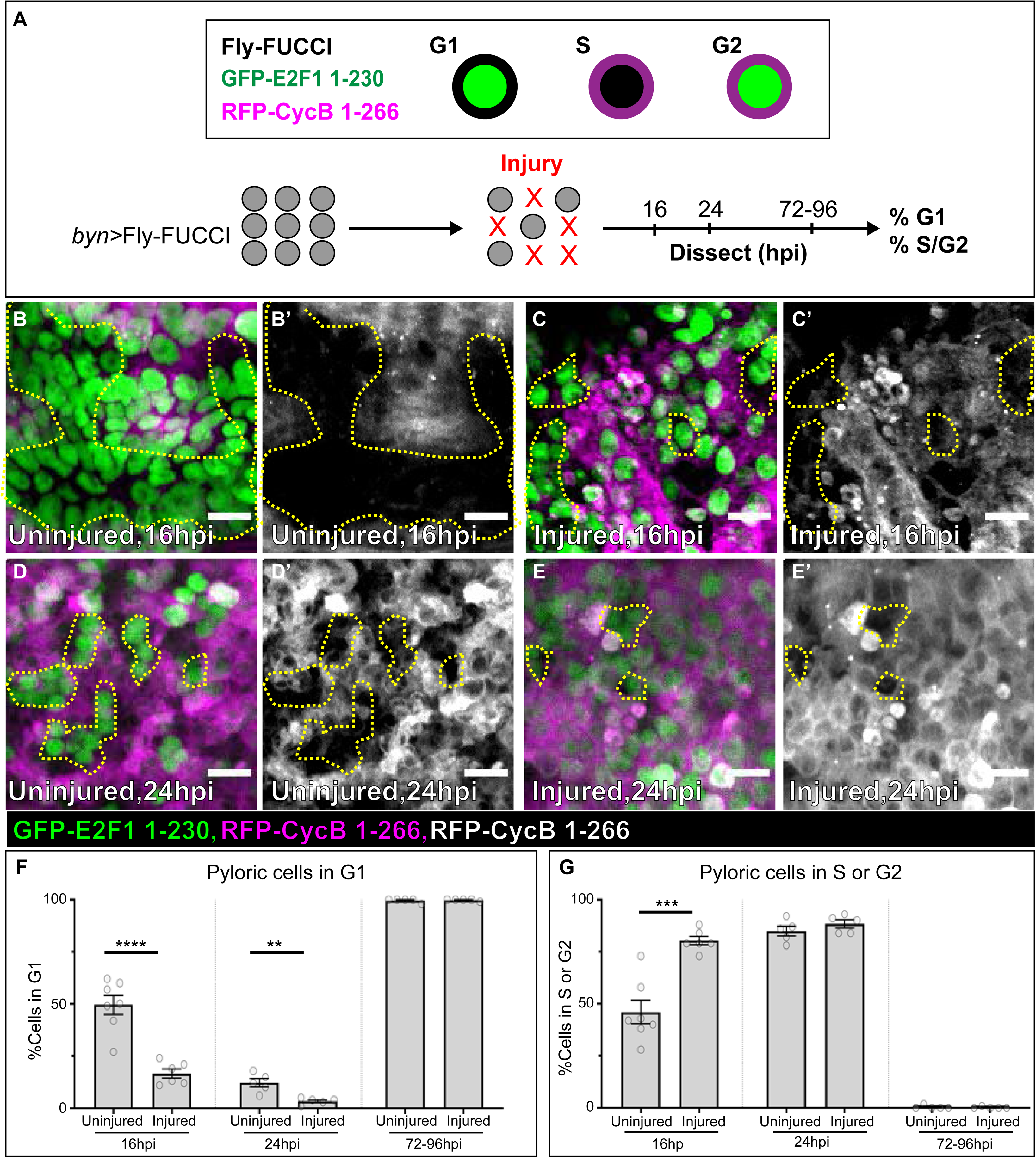
Pyloric cells accelerate G1 to allow for additional regenerative cell cycles. (**A**) Schematic illustrating the combination of DEMISE system with Fly-FUCCI system for analysis of cell cycle stages following injury. (**B-E’**) Insets of 16hpi (**B-C’**) or 24hpi (**D-E’**) developing pyloric cells with or without injury to wandering L3. E2F1.1-230 (green), CycB.1-266 (magenta or white). Yellow hashed line outlines cells at G1. (**F-G**) Quantification of (**F**) G1 and (**G**) S/G2 cells at different developmental times with or without injury at wandering L3 stage. Data represent mean ± SEM, N≥5 animals per condition, at least two replicates. Unpaired two-tailed t-tests. Scale bars 10µm.

### Fizzy-related activation by Ecdysone Receptor coincides with termination of the mitotic injury response

Given our finding that an L3W hindgut injury accelerates pyloric mitotic cell cycles to accomplish whole organ regeneration, we sought to identify the temporal signals that terminate this ability to respond to injury by mitosis. We previously found that in contrast to the mitotic cycles of the larval pylorus, adult pyloric cells instead undergo endocycles in response to tissue injury. Injury-induced endocycles in the adult pylorus increase cellular size (hypertrophy) and ploidy to restore tissue mass and genome content to the pre-injury state (Cohen et al., 2018). The switch to endocycles is governed by the Anaphase Promoting Complex/Cyclosome (APC/C) activator Fizzy-related (Fzr) (**Fig4A**, Cohen et al., 2018).

**Figure4.**
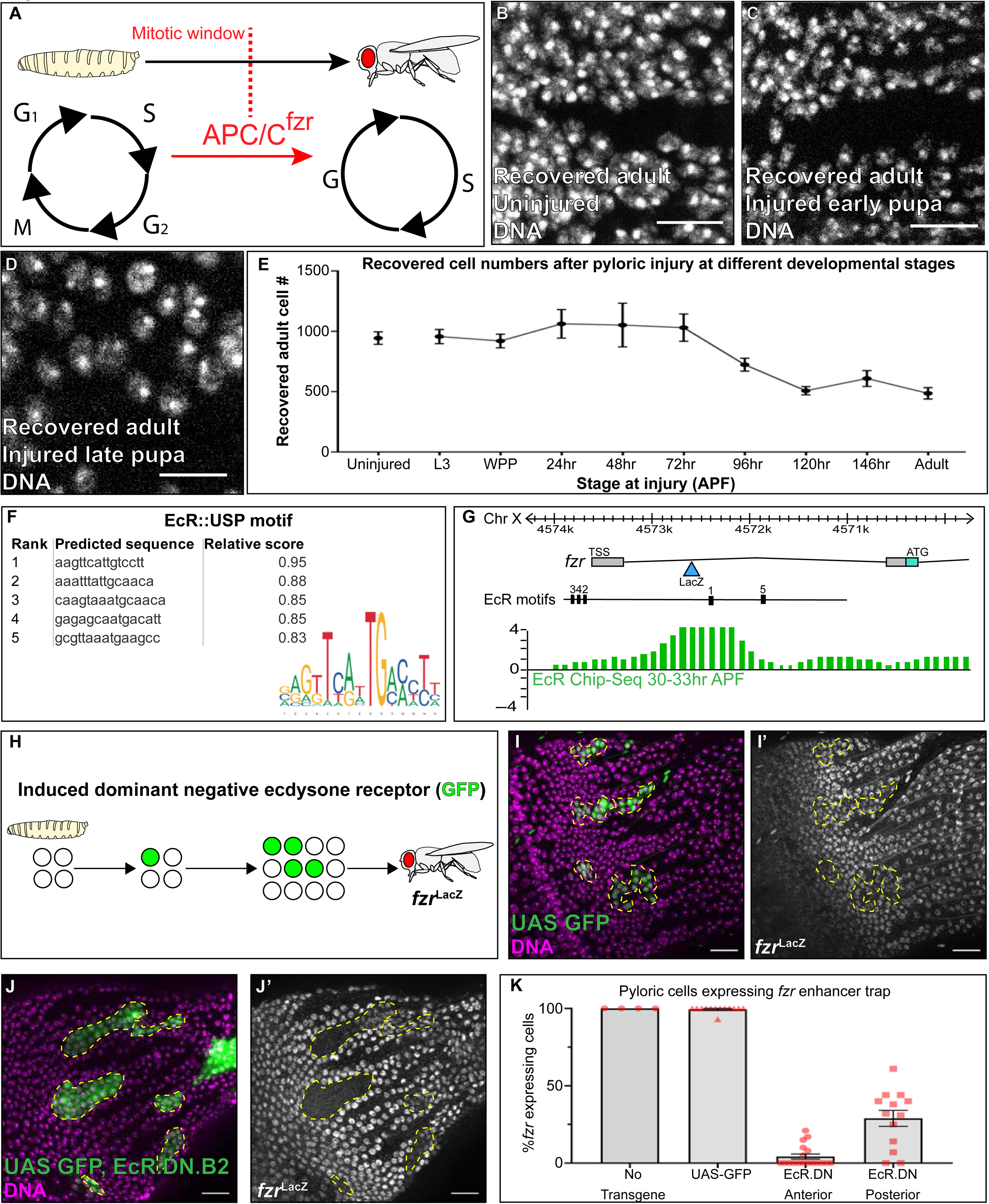
Ecdysone Receptor is required for *fizzy-related* enhancer activation during the termination of mitotic injury cell cycles. (**A**) Schematic of the injury response switch from mitosis at the wandering L3 stage to hypertrophy and endocycle at the adult stage. The switch is controlled by APC/C^fzr^ as previously published (Cohen et al., 2018). (**B-D**) Adult pyloric nuclei following (**B**) no injury, (**C**) injury induced 72 hours after pupation or (**D**) injury induced at 120 hours after pupation. Developmental stages represent time after pupation at 18C. Nuclei (DAPI, white). (**E**) Quantification of recovered adult cell number following injury at different developmental stages. Developmental stages represent time after pupa formation at 18C. Data represent mean ± SEM, N≥5 animals per condition with the exception of 24 and 48 hr APF where N=3 for each condition. At least two replicates. (**F**) Motif and relative scores of motif location in *fzr* enhancer region as obtained from JASPAR core. (**G**) modENCODE data identifying whole-animal EcR binding in *fzr* gene via ChIP-seq data. LacZ represents the location of the endogenous enhancer trap. TSS= transcriptional start site, ATG= translational start site. Thin lines= first and second introns, thick bars= exons (blue exon is retained in the coding sequence). Genomic locations are reported according to the modENCODE Genome Browser (**H**) Schematic of experimental design to induce hindgut-specific, dominant negative EcR clones throughout metamorphosis. (**I-J’**) Expression of a *fzr*^LacZ^ enhancer trap in adult pylori of **(I-I’**) GFP+ only clones or (**J-J’**) GFP+ Dominant negative EcR clones. Nuclei (DAPI, magenta), clones (GFP, green), *fzr*^LacZ^ (anti-Beta-Galactosidase, white). (**K**) Quantification of %*fzr^LacZ^* positive cells in adult clones expressing GFP or ecdysone receptor dominant negative in anterior or posterior pylorus. Data represent mean ± SEM, N≥5 animals per condition, at least two replicates. ANOVA, Tukey multiple comparisons test. Scale bars (**B-D**) 10µm, (**I-J’**) 20µm.

To pinpoint when injury-mediated pyloric cell mitoses are terminated during development, we first injured at several time points beginning at L3W, and examined recovered adults. To increase the temporal resolution of developmental progression, we raised and recovered animals at 18C, approximately doubling developmental time (**Methods**). To distinguish between injury-induced mitotic cycles and endocycles, we measured cell number of recovered adult pylori following injury at 9 distinct development times, over roughly one week of metamorphosis. Prior to mid-pupal stages, pyloric cells are capable of compensatory mitotic proliferation to restore correct adult hindgut cell number (**Fig4B-C, E**). The ability to undergo regenerative divisions prior to mid-pupation is consistent with our cell cycle marker analysis (**Figs1, 3**). Beginning at mid-pupation, cell number is abruptly no longer restored after injury and recovered nuclei appear larger, indicating that compensatory hypertrophy/endocycles begins at this point (**Fig4D-E, FigS2A**). We note that the end of injury-mediated mitotic cycles corresponds with the normal developmental termination of mitotic cell cycles, approximately 96 hours after pupa formation (APF) at 18C (**FigS1B**, 29C). To verify that the change in injury response coincides with the onset of endocycles, we measured cell ploidy before and after mid pupation. We observe an increase in ploidy of pylori injured after mid pupation (**FigS2A**). Together, our data support the existence of a short developmental window in which pyloric cells switch their injury response from that of compensatory mitotic divisions to compensatory endocycles and hypertrophy.

We next sought to identify developmental regulators of the injury response switch. We previously found that *fzr* drives the switch from mitotic cycles to endocycles in the pylorus, and that *fzr* expression is detectable in the adult (post-switch) pylorus, but not in the larva (pre-switch) (Cohen et al., 2018). We thus hypothesized that regulation of *fzr* expression underlies the pylorus injury cell cycle program switch. One such candidate for regulating developmental shifts at mid-pupation is the ecdysone steroid hormone. Ecdysone has been extensively studied as a regulator of developmental timing and progression (Riddiford et al., 2000). Recent studies in wing imaginal discs have also shown that signaling through the ecdysone receptor blocks regenerative capacity, leading to an inability to respond to injury following the prepupal hormonal pulse (Halme et al., 2010; Harris et al., 2016; Narbonne-Reveau and Maurange, 2019). However, unlike wing imaginal discs, we demonstrated that the hindgut does not lose the capacity to respond to injury following the prepupal ecdysone pulse, but rather changes its response to a *fzr-*dependent hypertrophic endocycles (**Fig4E**)(Cohen et al., 2018). We therefore explored whether ecdysone signaling regulates *fzr* activity in the hindgut, which would tie an animal developmental timing signal to a regulator of hindgut injury cell cycles.

We first identified a candidate *fzr* enhancer region based on the location of an established *lacZ* enhancer trap in the first intron of the *fzr* 5’ untranslated region. This trap is highly expressed in adult pylori (*fzr^G0418^,* Cohen *et al*. 2018). We then examined whether ecdysone regulates pyloric *fzr* expression during development. We analyzed potential motif sites in the *fzr* enhancer region and identified several strong Ecdysone Receptor (EcR) binding motifs (**Fig4F, Methods**). By analyzing modENCODE whole-animal ChIP-seq data (Celniker et al., 2009), we identified that EcR directly binds the *fzr* enhancer region at mid pupation (**Fig4G**). We next aimed to directly assess the effect of ecdysone signaling on hindgut development. Flies expressing a dominant-negative EcR in the hindgut throughout pupation die soon after eclosion due to gross defects in hindgut development and a lack of larval ileum histolysis (**FigS2B vs C**). Consequently, to ask whether ecdysone signaling is required for *fzr* enhancer expression in the adult pylorus, we induced hindgut-specific EcR dominant-negative clones (EcR.DN, **Fig4H, Methods**). We induced clones at the L3W stage to limit the effect of EcR.DN to metamorphosis. Flies expressing clonal EcR.DN in the hindgut show no morphological abnormalities and are viable. We then assayed expression of the *fzr lacZ* enhancer trap in adult animals containing either GFP+ EcR.DN clones, or GFP-only control clones (**Fig4I-K**). As previously observed, adult pyloric cells express *fzr lacZ* in the absence of injury (**Fig4I-I’**). Strikingly, pyloric clones expressing EcR.DN show a strong reduction in *fzr lacZ* expression (**Fig4J-J’**). The loss of this *fzr* enhancer trap expression is especially strong in clones of the anterior pylorus, where over 95% of cells show complete loss of *fzr lacZ* expression (**Fig4K**). Together, our results support a function for ecdysone signaling in regulating *fzr* activity in the developing hindgut. Additionally, the role of EcR in *fzr* enhancer regulation provides a direct link between steroid hormone signaling and termination of injury-mediated mitotic responses.

### *Dichaete* regulates the injury-mediated mitosis-to-endocycle switch

To further dissect a role for chromatin-level regulation in terminating injury-responsive mitosis, we closely examined *fzr* regulatory sequences. We sought to identify regions of the *fzr* enhancer near the EcR binding sites that are sufficient to reproduce the temporal and spatial hindgut expression of the *fzr lacZ* insertion. By analyzing modENCODE data, we confirmed that the region surrounding the EcR motif sites is positive for the active enhancer mark H3K27ac (**FigS3A**). To identify a specific *fzr* enhancer region that might be responsible for this hindgut activity, we cloned multiple fragments of the *fzr* sequence flanking various combinations of the five top motif sites for EcR binding (**Fig4F,G**, **Fig5A**) and generated transgenic lines. We note that flies injected with a fragment containing EcR motif sites 2-4 alone are not viable, whereas all other tested combinations (**Fig5A**) are viable. We then assessed the activity of these fragments to drive mCherry fluorescent protein in larval and adult guts (**Methods**). Of the three constructs, we identified one construct (*fzr*.B) that showed a pattern specific to the adult hindgut but not the larval hindgut (**Fig5B-C’**). The expression pattern of fragment *fzr.*B matches the temporal and spatial hindgut expression of the endogenous *fzr^LacZ^* enhancer trap (Cohen et al., 2018).

**Figure5.**
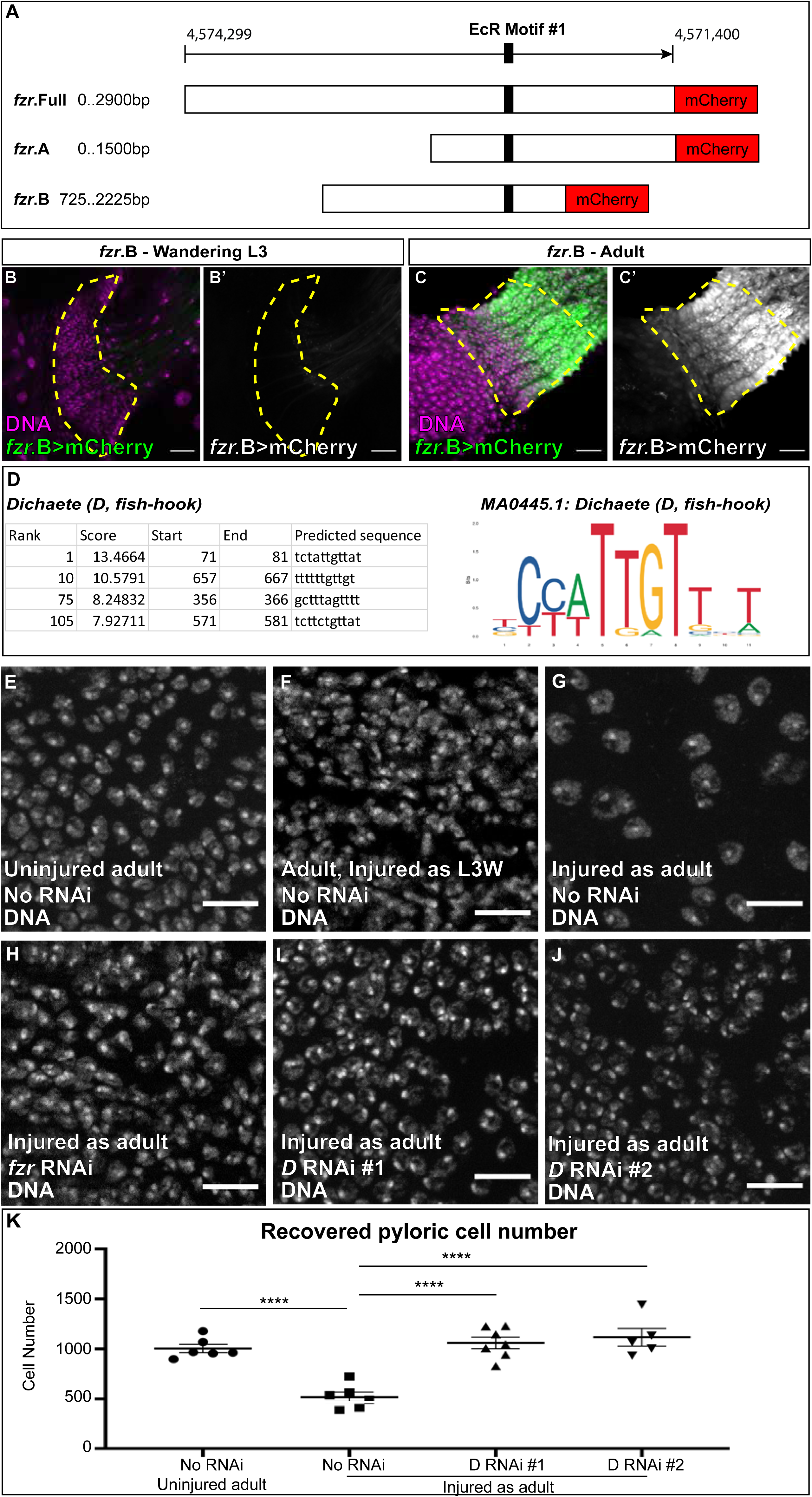
*Dichaete* regulates the hindgut injury mitosis-to-endocycle switch. (**A**) Schematic of cloned constructs. 1.5kb Fragments of the *fzr* enhancer were isolated and cloned in front of an mCherry coding sequence. Fragments were randomly integrated into the *Drosophila* genome and N≥2 lines of each fragment were assessed for expression. (**B-C’**) Expression of *fzr* enhancer *fzr*.B in (**B-B’**) the L3W pylorus and (**C-C’**) the adult pylorus. Nuclei (DAPI, white), mCherry (magenta or green). (**D**) JASPAR core motif analysis of non-overlapping region of *fzr*.B vs *fzr*.A identifies multiple potential binding sites for the *Dichaete* transcription factor motif. (**E-J**) Adult pyloric nuclei recovered after (**E**) no injury, (**F**) L3W injury (**G**) adult injury (**H**) adult injury in presence of *fzr* knockdown (**I-J**) adult injury in presence of *Dichaete* knockdown. Nuclei (DAPI, white). (**K**) Quantification of recovered cell number after adult pyloric injury in presence or absence of *Dichaete* RNAi. Data represent mean ± SEM, N≥5 animals per condition, at least two replicates. ANOVA, Tukey multiple comparisons test. Scale bars (**B-C’**) 20µm, (**E-J**) 10µm.

Fragment *fzr.*B contains the highest ranked EcR motif site (**site #1, Fig4F, G, Fig5A**), adjacent to both the *lacZ* trap and the modENCODE pupal EcR Chip-Seq peak. However, all three constructs (**Fig5A**) contain this highest ranked EcR binding motif site, yet not all drive mCherry expression in the hindgut (**Fig5B**, **FigS3B-E’**). Rather, *fzr*.Full and *fzr*.A fragments show differential temporal and spatial expression patterns, driving mCherry expression in the Malpighian tubules and midgut enterocytes respectively (**FigS3B-E’).** The differential expression of the constructs suggests that this strong EcR binding site is not sufficient to induce *fzr* enhancer fragment expression, or that other sequences present in *fzr*.Full and *fzr*.A fragments but not in *fzr.*B might include repressors of *fzr* activation. These results align with previous observations in wing imaginal discs, where EcR is responsible for changing chromatin architecture rather than directly activating genes (Ma et al., 2019; Uyehara et al., 2017).

Given these findings, we next sought to identify transcription factors that might control the injury response switch. To identify relevant candidates, we analyzed the adult hindgut-expressed *fzr* enhancer fragment for transcription factor binding sites. To limit the scope of the sequence to potential hindgut activators of *fzr,* we analyzed the unique 725bp region of *fzr*.B compared to the non-hindgut expressing *fzr*.A. From motif analysis, we identified two strong motif sites of the Sox-domain-containing transcription factor *Dichaete (D, fish-hook)* (**Fig5D, Methods**). *Dichaete* has previously been implicated in embryo segmentation, cell fate, and differentiation (Ma et al., 1998; Russell et al., 1996; Zhao and Skeath, 2002) Additionally, *Dichaete* has previously been described to express in all larval hindgut cells as well as multiple types of imaginal discs, suggesting a role for the gene at metamorphosis (Mukherjee et al., 2000). However, the role of *Dichaete* in injury responses remains unexplored.

To test whether *Dichaete* is a regulator of the hindgut developmental injury response switch, we utilized our DEMISE system to injure the adult gut in the presence or absence of *Dichaete. Dichaete* knockdown in the hindgut does not lead to morphological defects or cell fate changes in the absence of injury, as assayed by the maintained expression of hindgut-specific *byn*>*gal4* (**FigS3F**). We then injured adult hindguts of wild-type animals or animals expressing *Dichaete* RNAi throughout metamorphosis. We confirmed injury to the hindgut pylorus by DCP1 staining immediately following injury (**FigS3G**). Once we established that injury occurs, we allowed flies to recover and measured pyloric cell number as done previously (Cohen et al., 2018) to assess if injury is compensated by mitosis or endocycle. As we previously established, in contrast to the mitotic response following L3W injury, adult wild-type flies respond to injury without mitosis and instead increase size/ploidy of remaining nuclei (**Fig5E-G**). Loss of *fzr* in the adult pylorus enables cell number restoration without ploidy increase after adult injury, as previously described (**Fig5H,** Cohen et al., 2018). Using two separate RNAi constructs, *Dichaete* RNAi in the adult pylorus also restores cell number following acute injury, without a noticeable nuclear size increase (**Fig5I-K**). These results implicate *Dichaete* as a regulator of the pyloric injury switch from mitotic cycles to endocycles. Thus, our analysis of a hindgut *fzr* enhancer fragment identified the *Dichaete* transcription factor as a negative regulator of injury-induced mitotic cell cycles.

## Discussion

### Accelerating the cell cycle as a mechanism to regenerate organs within developmental time constraints

The connection between organism development and the capacity to undergo regeneration has been observed in diverse tissues and organisms (reviewed in Poss 2010; Seifert and Voss 2013; Yun 2015). Here, we use the *Drosophila* hindgut pylorus as a model to understand how regenerative mitotic cell cycles can be coordinated with tissue development. We show that a delay in development is not essential for hindgut regenerative mitotic cell cycles. Rather, we show that injury to the pylorus of wandering 3^rd^ instar larvae shortens G1 phase to accelerate the mitotic cell cycle. This allows for increased regenerative cell divisions within a developmental time constraint. We further identify molecular signals that terminate this mitotic regenerative capacity in the developing pylorus. Our work reveals that the steroid hormone ecdysone receptor and the Sox family transcription factor *Dichaete* switch the cellular injury response from relying on mitotic cycles to relying on endocycles and hypertrophy. Our data reveal a mechanism by which flexible mitotic cell cycle dynamics during a succinct period of organ development enable regeneration (**Fig6**).

**Figure6.**
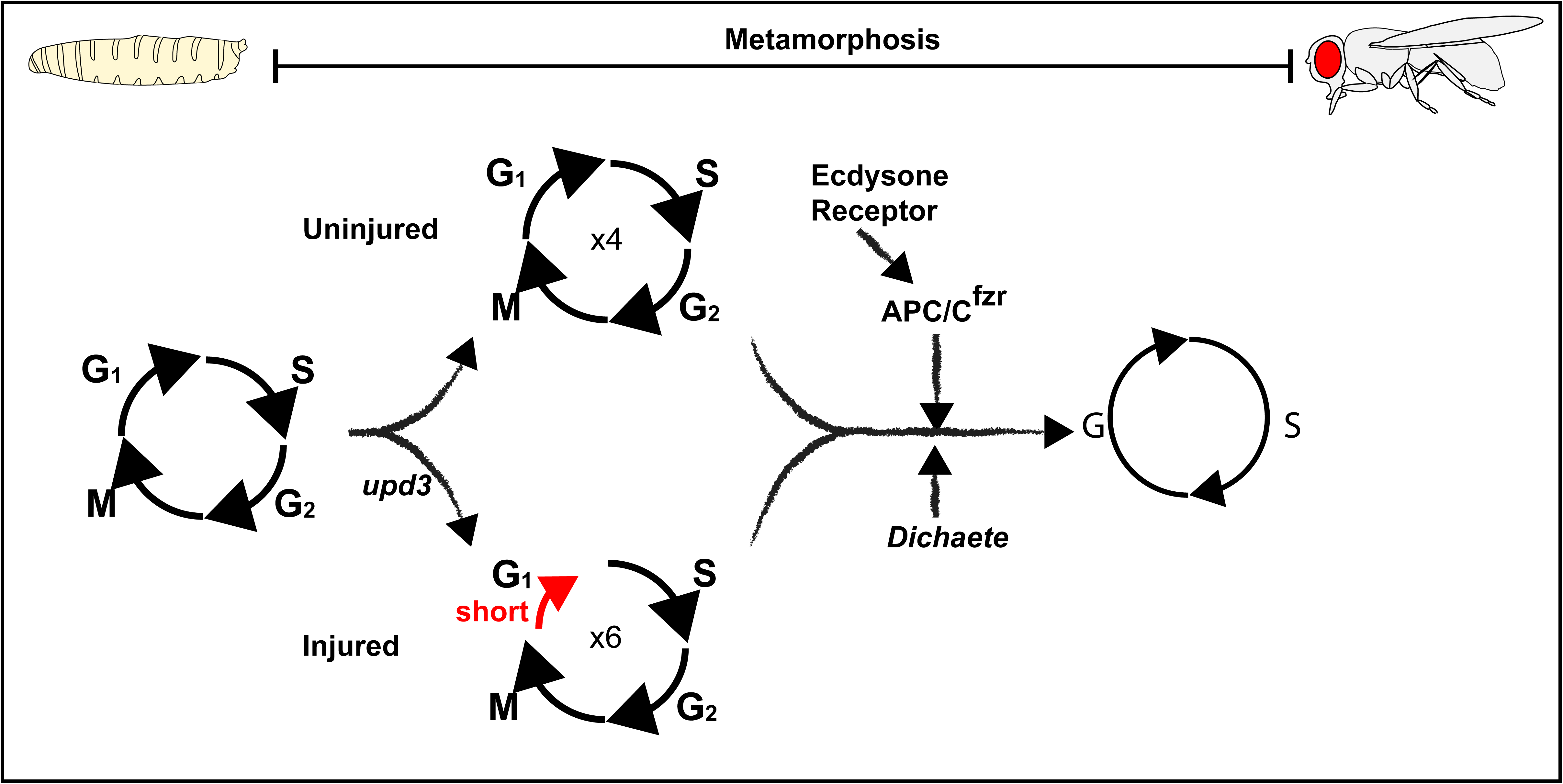
Model: Regeneration under developmental time constraints. Injury to the L3W stage does not delay development. Injured pylori undergo 2 additional mitotic cycles in order to restore injured cell number. Additional mitotic cycles occur through decreased time spent in G1 and require the Upd3 JAK/STAT cytokine. The accelerated divisions are terminated by the ecdysone receptor, which activates an *APC/C^fzr^* enhancer, as well as the SOX transcription factor *Dichaete*.

The ability of the hindgut to undergo mitotic regeneration without delaying development is distinct from that of the *Drosophila* wing disc, which cannot regenerate after pupation onset (Halme et al., 2010). Our work suggests that while a developmental delay can be utilized to prolong a transient mitotic regenerative state, it is not essential for regenerative divisions in other contexts. These findings show that hindgut mitotic regeneration under time constraints requires cell cycle acceleration. To accomplish such acceleration, our work here reveals that G1 regulation is likely a critical node of regulation. G1 is a time of extrinsic sensing before passing through start of the next cycle. Our data suggest that injury-responsive JAK/STAT signals, mediated by the cytokine Upd3, may provide an injury-based extrinsic cue to progress quickly through the G1/S transition to accomplish mitotic organ regeneration. Further, our findings using heterozygous *upd3* animals highlight that precise cytokine levels are critical for injury-induced cell cycle acceleration. We note that previous studies in the *Drosophila* eye and wing disc found that misexpression of the JAK/STAT cytokine *upd* also leads to cell cycle acceleration (Bach et al., 2003; Rodrigues et al., 2012). We also note that acceleration of mitotic cycles under stress conditions have been identified in other stem cell/regenerative contexts, including regeneration of the axolotl spinal cord (Rost et al., 2016), as well as in mammalian hematopoietic progenitors, where shortened G1 may also be involved (Guo et al., 2014; Mende et al., 2015). Future work in the pylorus can provide an accessible model of conserved cell cycle control in organ regeneration.

### A role for hormonal and transcriptional regulation in coordinating injury cell cycles

Our results identify both hormonal and transcriptional cues that temporally alter tissue injury cell cycles. We previously identified the mitotic inhibitor APC/C^fzr^ as a regulator of injury responsive cell cycle alterations in the pylorus (Cohen et al., 2018). In the adult pylorus, *fzr* upregulation leads to degradation of mitotic machinery prior to injury, priming the tissue towards a hypertrophic injury response (Cohen et al., 2018). We now identify that the ecdysone steroid hormone receptor activates a *fzr* enhancer to terminate injury-mediated mitosis. Regulation of this *fzr* enhancer by the ecdysone receptor directly links systemic hormonal factors to an injury-mediated mitosis to endocycle switch.

Our data supports an emerging concept in the literature, where developmentally timed systemic signals impact tissue injury responses. Similar to the pylorus, the ecdysone steroid hormone limits the regenerative capacity of the *Drosophila* wing imaginal disc (Narbonne-Reveau and Maurange, 2019). In response to injury, retinoic acid and insulin-like peptide 8 are released from the wing disc to delay ecdysone production and organismal development (Halme et al., 2010). In mammals, studies have identified a role for circulating factors in maintenance or loss of regenerative capacity (Avci et al., 2012; Conboy et al., 2005; Elabd et al., 2014; Hirose et al., 2019; Rebo et al., 2016). Notably, circulating hormones such as oxytocin and thyroid hormone play a role in regulation of transient regenerative capacity (Avci et al., 2012; Elabd et al., 2014; Hirose et al., 2019). Our model supports the importance of hormonal signals in the control of injury cell cycle states.

By further exploring the connection between EcR binding and *fzr* enhancer, we identified a hindgut hindgut-specific *fzr* enhancer fragment, which then led us to identify an additional regulator of the injury response switch: the Sox-domain-containing transcription factor *Dichaete*. Sox transcription factors have been involved in regeneration in other species including zebrafish spinal cord regeneration (Guo et al., 2011) and mouse nerve regeneration (Jing et al., 2012). Whether *Dichaete* acts as a direct activator of *fzr* or performs its function indirectly through interaction with regulatory pathways or chromatin is a question for future study.

Our ability to identify novel regulators of regeneration capacity through analysis of enhancer elements emphasizes the role of chromatin regulation in injury-responsive cell cycle programming. Indeed, tissue regenerative enhancer elements have been identified in both vertebrates and invertebrate species. In the *Drosophila* wing disc, activation of a *wingless* damage-responsive enhancer fragment is required for regeneration following injury (Harris et al., 2016; Smith-Bolton et al., 2009). As development progresses, this *wingless* enhancer is silenced by an adjacent element, supporting a role for chromatin environment in both activating and silencing regeneration (Harris et al., 2016). Further research in *Drosophila* wing discs has identified ecdysone-induced factors to change genome wide chromatin accessibility around cell cycle genes (Ma et al., 2019; Uyehara et al., 2017). Tissue regeneration enhancer elements (TREE) have also been identified in zebrafish, as well as acoels, further emphasizing the evolutionary connection between enhancer activation and regenerative capacity (Gehrke et al., 2019; Kang et al., 2016). Our finding of a *fzr* enhancer fragment that is both spatially and temporally upregulated in the adult hindgut to terminate injury mitotic cycles supports a role for chromatin accessibility regulation upon tissue injury.

In summary, this study highlights the *Drosophila* pylorus as a novel model for studying how mitotic regeneration can be coordinated and achieved within developmental tissue programming. Future work in this system will illuminate how systemic signals and chromatin changes can regulate cell cycle dynamics to control tissue injury responses.

## Acknowledgements

The following kindly provided reagents used in this study: Bloomington Drosophila Stock Center, Developmental Studies Hybridoma Bank, Vienna Drosophila Resource Center. We thank Bernard Mathey-Prevot, Stefano Di Talia, and Ruth A. Montague for comments on the manuscript. This project was supported by NIGMS grant GM118447 to DF.

## Author contributions

Conceptualization: E.C., D.T.F.; Methodology: E.C.; Validation: E.C.; Formal Analysis: E.C.; D.T.F.; Investigation: E.C.; Data curation: E.C.; Writing - original draft: E.C., D.T.F.; Visualization: E.C.; Supervision: D.T.F.; Funding acquisition: D.T.F.

## Declaration of Interests

The authors declare no competing interests.

## Materials and Methods

### Fly stocks and *Drosophila* genetics

Full genotypes are described at flybase.org. Flies were raised on standard *Drosophila* media (Archon Scientific, Durham) at 25C unless reported otherwise. All adult dissections were performed 4-7 days following eclosion. With the exception of *upd3*Δ hemizygous males, data were collected and quantified from female flies. *upd*3Δ homozygous virgin females were used for all crosses to avoid rescue by maternal load. The following publicly available stocks were used in the study and their Bloomington *Drosophila* Stock Center number (BS#) provided: *fzr^G0418^* (#BS 12297), *hsFLP12;Sco/CyO* (#BS 1929), *UAS-EcR.B1-ΔC655.F645A* (#BS 6869), *upd3Δ* (#BS 55728), UAS-*D.*RNAi#1^TRiP.JF02115^ (#BS 26217), UAS-*D.*RNAi#2 ^TRiP.HMS01150^ (#BS 34672), *ptub-FRT-Gal80-FRT* (#BS 38881). Additionally, the following flies were used in the study: *byn>Gal4, UAS-hid, UAS-reaper* (Cohen et al., 2018; Fox and Spradling, 2009; Sawyer et al., 2017). The DEMISE component *UAS-FRT-stop-FRT*-*rpr* was previously generated in our lab and is available upon request (Cohen et al., 2018).

All UAS transgenes were induced by *byn>Gal4.* Unless indicated, all injury protocols were performed as previously described using the DEMISE injury system (*hsFLP;UAS-FRT-stop-FRT-rpr#10-3*,Cohen *et al*. 2018). For wandering L3 (L3W) injury, flies were kept at 29C throughout development and collected at the wandering L3 stage. Larvae were then subjected to a 35 min 37C heat shock, a sub-lethal dose of injury. Flies were then shifted back to 29C to allow continued expression of transgenes and apoptotic genes. Injury for the following experiments: developmental delay (**Fig1B-D**), pupal injury (**Fig4E**) and morphology following injury of *upd3*Δ animals (**Fig2A-C**) was performed by regulating UAS-*hid* and UAS-*rpr* expression using the Gal80^ts^ repressor as previously described (Cohen et al., 2018). In these experiments, flies were kept at 18C until the desired developmental stage and shifted to 29C for 16 hours. Flies were then transferred to 18C for measuring development or adult dissection 4-7 days following eclosion.

Ecdysone receptor dominant negative clones were induced by flipping out a ptub-FRT-Gal80-FRT repressive cassette (Bohm et al., 2010). Flies containing (*hsFLP/fzr^G0418^; EcR.DN; byn>Gal4, UAS-NLSGFP, ptub-FRT-Gal80-FRT*) were kept at 29C and subjected to a 30-minute heat shock at 37C at the wandering L3 stage. Induction of FLP by heat shock leads to removal of Gal80 cassette and stochastic expression of transgene of interest by *byn >Gal4* throughout metamorphosis. Flies were then shifted back to 29C and dissected 4-7 days post eclosion.

### Enhancer fragment cloning and motif analysis

Identification of H3K27ac and ecdysone receptor potential binding sites was based on publicly available data from modENCODE (Celniker et al., 2009). Motif analysis was performed using the JASPAR core Scan analysis tool (Fornes et al., 2020). The top 5 ranking non-overlapping motifs are reported. Candidate *fzr* enhancer fragments were isolated by PCR from *w^1118^* flies, sequenced and compared to a reference genome. The fragments were then inserted into the Gateway entry vector pDONOR221 (ThermoFisher Scientific) using Invitrogen Gateway BP Clonase II Enzyme Mix. Fragments were then inserted into a publicly available destination vector pHPdestmCherry (Addgene #24567, Boy et al. 2010), using the Invitrogen LR Clonase Enzyme Mix (ThermoFisher Scientific). The final construct was integrated at random genomic sites, and two homozygous viable lines were selected per fragment. All flies and constructs are available upon request.

### Ploidy measurements and Staining

For ploidy measurements, guts were dissected in 1X PBS, prepared and measured as previously described (Cohen et al., 2018; Fox et al., 2010; Losick et al., 2013). Adult pyloric ploidy represents an average of N>100 cells per animal, normalized against haploid cells in the testis. For ploidy calculations, images were obtained with an upright Zeiss AxioImager M.2. For all other experiments, dissection, fixation and staining protocols were performed as previously described (Cohen et al., 2018; Sawyer et al., 2017). The following antibodies were used in this study: Beta-Galactosidase (Abcam, ab9361, 1:1000), DCP1 (Cell Signaling, Asp261, 1:500), Phospho-Histone H3 (Cell Signaling, #9706, 1:1000). Secondary antibodies were Alexa Fluor dyes (Invitrogen, 1:500). Tissues were mounted in Vectashield (Vector Laboratories Inc.). Images were obtained with an upright Zeiss AxioImager with Apotome.2 processing, inverted Leica SP5 or Andor Dragonfly Spinning Disk Confocal. Image analysis was performed using ImageJ (Schneider et al., 2012), including adjusting brightness/contract, Z projections, cell counts, and integrated density quantification. Image stitching (**Fig2A, FigS2B-C**) was performed using ImageJ grid/collection stitching plugin (Schneider et al., 2012).

### Statistical analysis and reporting

Statistical analysis was performed using GraphPad Prism 8.2.0. Statistical tests are detailed in figure legends. P and adjusted P value reporting is as follows: (p>0.05,not significant); (p<0.05,*); (p<0.01,**); (p<0.001,***); (p<0.0001, ****).

## Supplementary Material

**FigureS1.**
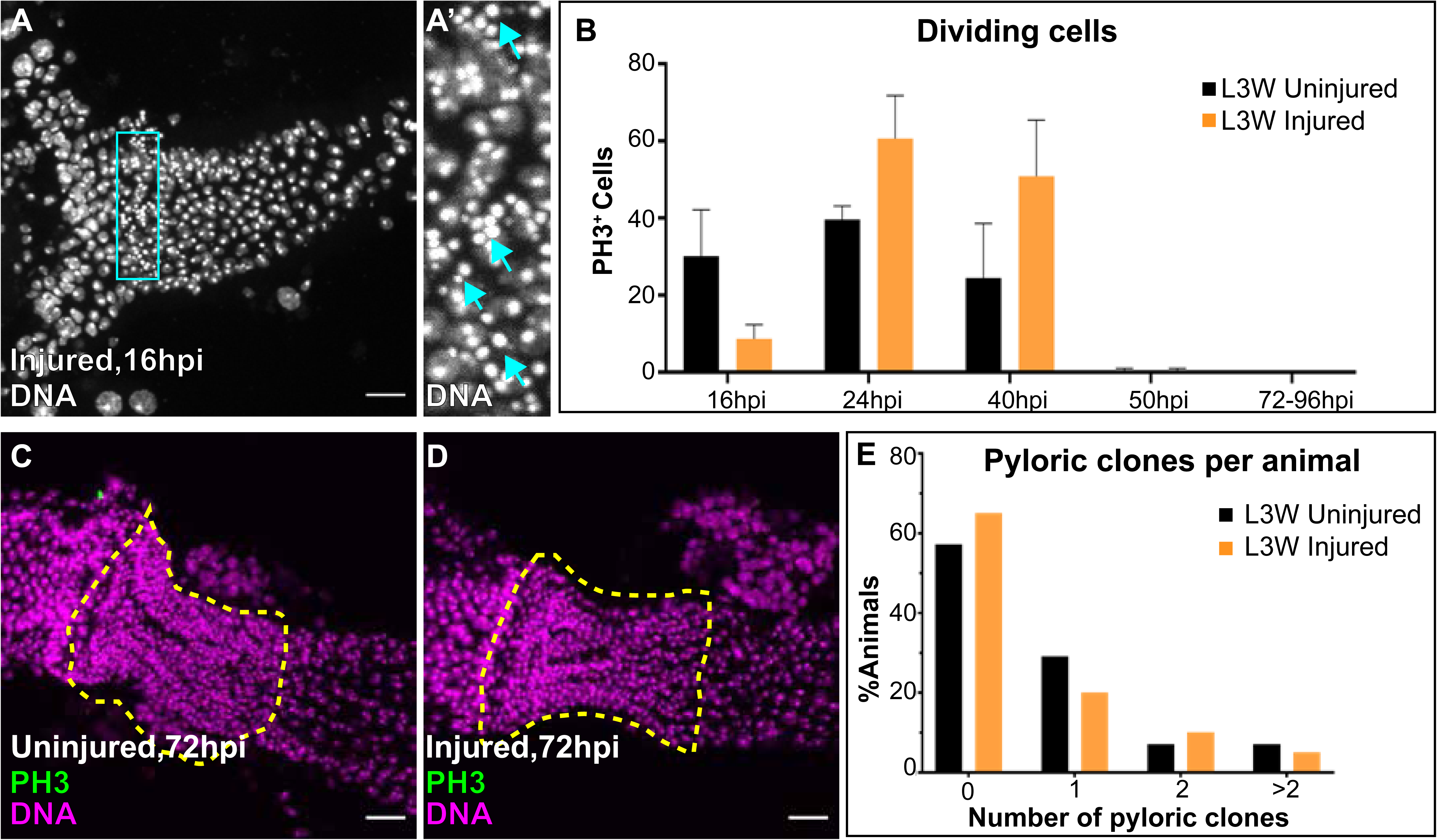
Supplement to Figure1. (**A-A’)** Pyknotic nuclei at 16hpi. Nuclei (DAPI, White), blue outline represents the location of the (**A’**) inset. Blue arrows point to pyknotic nuclei (**B**) Quantification of raw number of mitotic nuclei in developing pylorus. Data represent mean ± SEM, N≥5 animals per condition with the exception of 50hpi where N=4 (uninjured) and N=2 (injured). All data represents at least two replicates. (**C-D**) Lack of mitotic figures at 72hpi. Phospho-HistoneH3 (green), and nuclei (DAPI, magenta) are marked. Yellow hash outlines the pylorus, derived by *byn* expression and hindgut morphology. (**E**) Number of adult pyloric clones per animal in presence or absence of injury under sparse clonal induction at the L3W stage. Scale bars 20µm.

**FigureS2.**
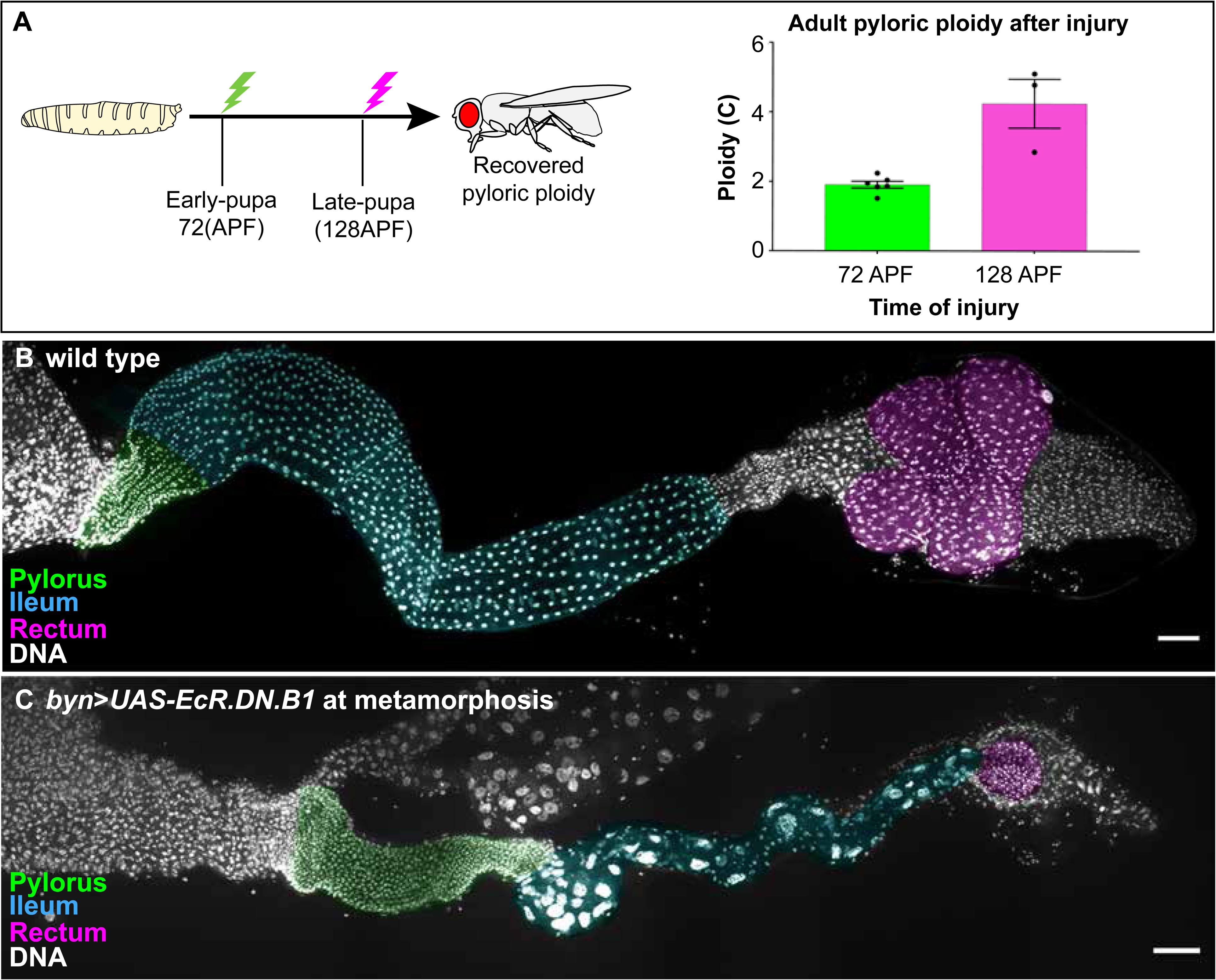
Supplement to Figure4. (**A**) Ploidy quantifications of recovered adult pylori following injury at early pupa (Green, 72hr, 18C) and late pupa (Magenta, 124hr, 18C). Data represent mean ± SEM, N≥3 animals per condition and marked by symbols on graph. recovered cells counted>100 per animal. All data represents at least two replicates. Unpaired two-tailed t-test. Lightning bolt illustrations represent injury induction. (**B-C**) Pharate hindgut morphology in (**B**) wild-type and (**C**) animals expressing hindgut-specific dominant negative ecdysone receptor throughout metamorphosis. Nuclei (DAPI, white) and false color identifies the tissues of the adult hindgut: Pylorus (green), Ileum (cyan) and Rectum (magenta). Stitching was performed to represent the entire hindgut in a single image. Scale bars 50µm.

**FigureS3.**
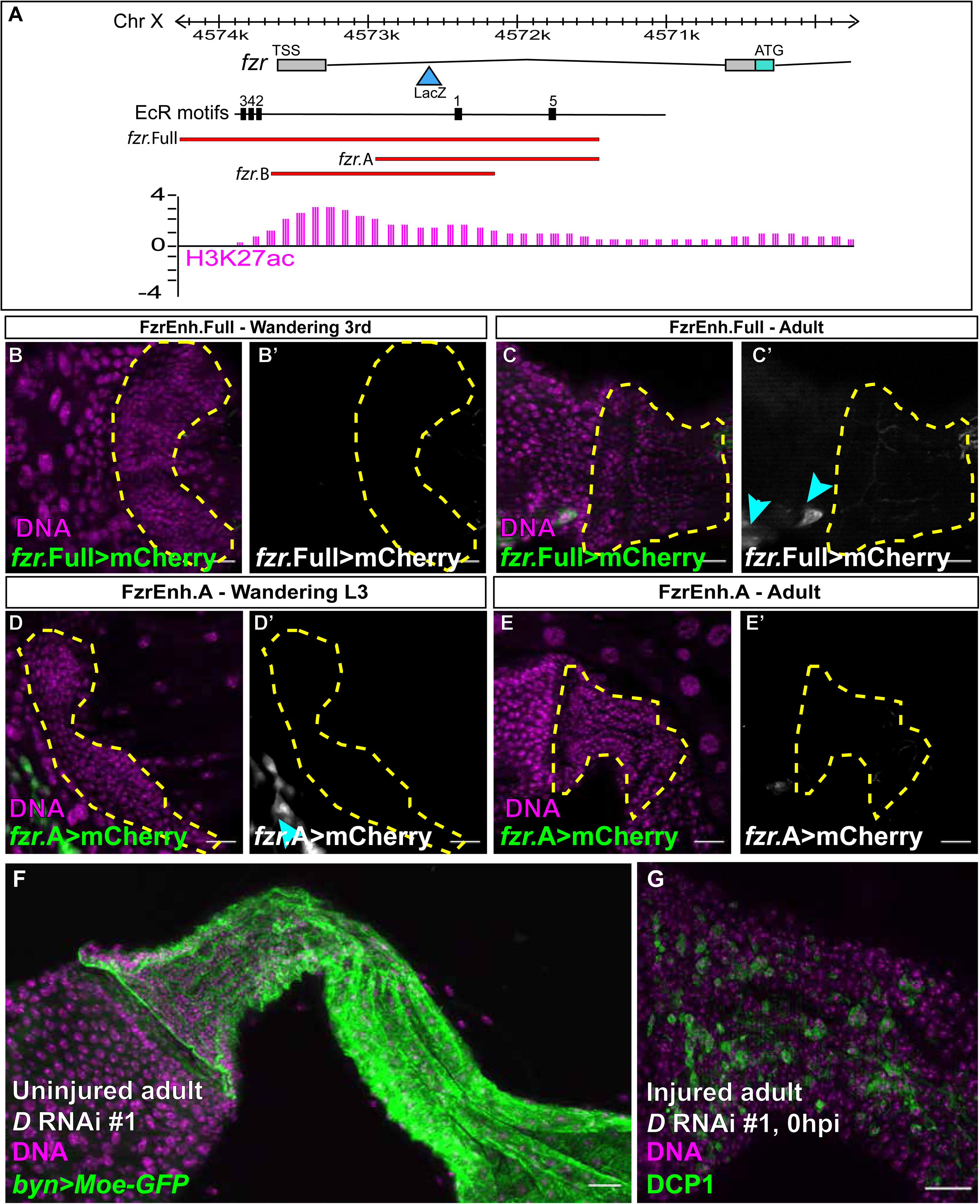
Supplement to Figure5. (**A**) A candidate *fzr* enhancer region as identified by H3K27Ac marks in modENCODE overlaps with ecdysone motifs. Cloned fragments are overlaid on genomic map in red (**B-C’**) Expression of *fzr* enhancer *fzr*.Full in (**B-B’**) the L3W pylorus and (**C-C’**) the adult pylorus. Nuclei (DAPI, white), mCherry (magenta or green). Cyan arrows point to Malpighian tubule expression. (**D-E’**) Expression of *fzr* enhancer *fzr*.A in (**D-D’**) larval pylorus and (**E-E’**) adult pylorus. Nuclei (DAPI, white), mCherry (magenta or green). Arrows point to midgut enterocyte expression. (**F**) *byn* expression in adult hindguts expressing *Dichaete* knockdown throughout metamorphosis. (**G**) DCP1 staining of injured adult hindguts expressing *Dichaete* RNAi throughout metamorphosis. Nuclei (DAPI, magenta), apoptosis (DCP1, green). Scale bars 20µm.

